# Mouse primary visual cortex neurons respond to the illusory “darker than black” in neon color spreading

**DOI:** 10.1101/2022.07.24.501311

**Authors:** Alireza Saeedi, Kun Wang, Ghazaleh Nikpourian, Andreas Bartels, Nelson K. Totah, Nikos K. Logothetis, Masataka Watanabe

## Abstract

Illusions are a powerful tool for studying the single neuron correlates of perception. Here, we introduce the neon color spreading (NCS) illusion in mice and report the neuronal correlates of illusory brightness, which has heretofore only been studied using human fMRI. We designed a novel NCS paradigm to evoke the percept of an illusory drifting grating and analyzed the activity of 520 single units in the mouse primary visual cortex (V1). A substantial proportion of V1 single units (60.5%) responded to illusory gratings with direction tuning matched to their preferred direction, which was determined using physically presented luminance-defined gratings (LDG). Moreover, by presenting LDG gratings with a 180° phase shift relative to NCs gratings, we show that spatial phase tuning shifted 180° for most single units. This finding conclusively demonstrates that V1 single units respond to illusory brightness. Using this novel mouse paradigm, we show that responses to illusory gratings have a lower magnitude and are delayed relative to physical gratings. We determined where V1 single units fell in the V1 cellular hierarchy (based on their susceptibility to surround suppression, their putative classification as interneuron or pyramidal neuron, and designation as a simple or complex cell) and found that higher-level V1 single units are more responsive to NCS stimuli. These findings resolve the debate of whether V1 is involved in illusory brightness processing and reveal a V1 hierarchical organization in which higher-level neurons are pivotal to the processing of illusory qualities, such as brightness.

## Introduction

Non-human primate and cat V1 single cells respond to physical brightness^1-6^, but their response to illusory brightness is unexplored^7,8^. Moreover, fMRI studies in humans have been inconclusive, with some studies reporting a correlation between V1 BOLD signal and perception of illusory brightness^9-11^, while others found no such correlation^12-14^. Thus, the V1 neuronal response to illusory brightness is uncertain in humans and animals^15-18^.

In the present study, using *in vivo* electrophysiology in mouse VI, we probe the neural correlates of neon color spreading (NCS) ^19,20^, an illusion which has been shown in humans to combine different perceptual qualities, such as filling-in and perception of illusory contours and brightness^21^. We show that single units recorded in mouse V1 respond to NCS stimuli designed to evoke the percept of an illusory drifting grating. Neuronal responses were compared with a physical drifting grating that was 180° phase shifted relative to the illusory grating. By analyzing the spatial phase tuning properties of single units this approach allowed us to demonstrate that V1 single units respond to illusory brightness.

Using this mouse paradigm, we were able to probe the putative neuronal connectivity of the intra-Vl cellular hierarchy^22-26^ involved in illusory brightness processing by studying the relationship between single unit responses to the illusory stimuli and surround modulation^27,28^, complex-simple cell modulation^29-32^, as well as by characterizing units as putative inhibitory interneurons and putative excitatory pyramidal neurons ^33,34^. Our results show that single units with a stronger preference for NCS stimuli are at a higher level in the functional or physiological hierarchy of mouse V1.

## Results

Wc recorded 636 V1 single units in 6 head-fixed mice passively viewing visual stimuli (Fig. 1). Before the experiment, we made two preliminary recordings aimed at locating single unit receptive fields and characterizing size tuning. Briefly, we first performed receptive field mapping to estimate the center of the receptive field using the multi-unit response to black squares (15° widths) presented on a gray background at locations selected in a pseudo-random order from an 8 by 13 grid. Next, we recorded unit responses to circular patches of drifting gratings with different sizes (2.5° - 45°) presented in the receptive field center. These recordings were subsequently used to characterize size tuning. Following these preliminary recordings, we began recording the neuronal response to the stimuli shown in Fig. 1 to study whether V1 single units respond to illusory brightness.

**Fig. 1:**
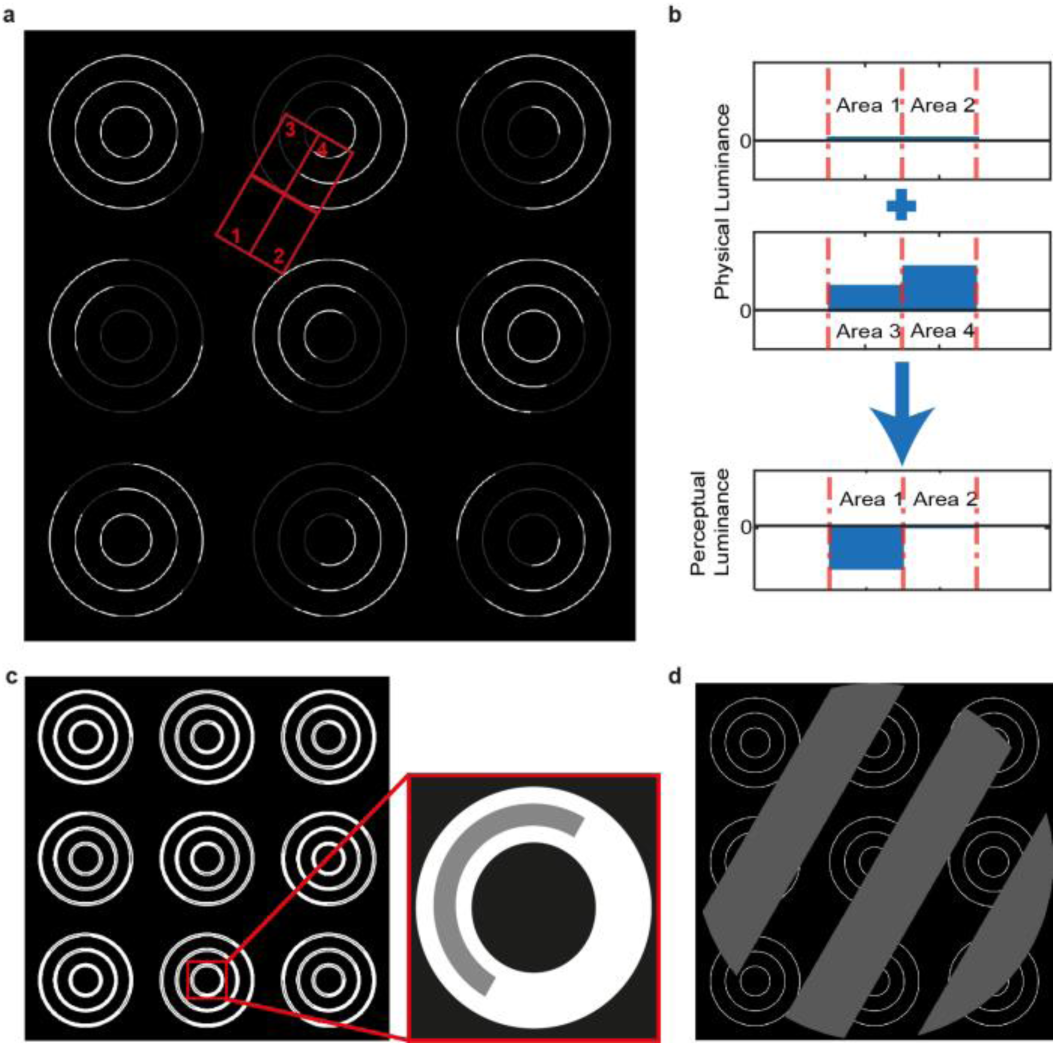
The visual stimuli consisted of NCS stimuli that evoked illusory luminance to form a drifting grating and control stimuli that either blocked the illusion or presented a physical grating. **a**, An example of the achromatic NCS stimulus presented to mice. The stimulus consists of 9 sets of white concentric circles. Each set of concentric circles contains gray segments at different positions. As can be seen, the gray color is diffusing into the empty area between concentric circles. This generates an illusory grating that appears darker than the surrounding black background. Changing the location of the grey segments on the concentric circles will change the orientation of this illusory grating. **b**, The schematic shows the physical and perceptual luminance in the four arbitrary areas bounded by red boxes and marked as areas 1,2,3, and 4 in panel a. Luminance differences in areas 3 and 4 make an illusory perceptual luminance difference between areas 1 and 2 due to the diffusion of the gray color from area 3 to area 1. **c**, In the diffusion-blocked control (DBC) stimulus condition, the concentric circles are constrained by an outer white ‘band’, which serves to block the diffusion of gray color while maintaining the presentation of the physical stimulus (i.e., the concentric circles). **d**, A control stimulus that provided a physical luminance-defined grating (LDG) was used to compare an actual grating with the perceptual grating evoked by the NCS illusion in panel a. This grating was presented over a background compound of a steady concentric circle with a temporal and spatial frequency identical to the illusory grating generated by the NCS stimulus.

We presented three types of stimuli centered on the receptive field and covering 35° width. The first stimulus type was the NCS stimulus (Fig. la). This stimulus consisted of an array of white concentric circles presented on a black background. Each of the concentric circles in the array contains grey segments at different positions. These grey segments are aligned in a way that diffusion of gray color into the background produces an illusory grating which seems darker than the surrounding black background. This NCS stimulus is an achromatic version of the NCS illusion in human studies^21^. Depending on which segments of the concentric circles are grey, the orientation of the grating changes. Grey segments were introduced at each time frame to generate a drifting illusory grating (Supplementary Movie 1).

The second and third types of stimuli were control conditions. The diffusion-blocked control (DBC) stimulus (Fig. 1c) had identical temporal dynamics to the NCS stimulus. However, each concentric circle was constrained by two static circles, which led to the extinction of the illusory percept by disrupting the diffusion of brightness (Supplementary Movie 2). The DBC stimulus was used to verify that a V1 response to the NCS stimulus was due to the processing of an illusory grating, as opposed to the physical stimulus changes within the receptive field. The other control stimulus was a luminance-defined grating (LDG), which was a drifting grating presented in the foreground over the concentric circles (Fig. 1d, Supplementary Movie 3). The stimulus had the same spatial and temporal frequency as the illusory grating generated by the NCS stimulus. This stimulus served to provide a control condition in which a physical drifting grating was presented in order to compare tuning properties of single units for illusory gratings (NCS stimuli) with physical gratings. The same gray color was used in all conditions to have comparable neuronal responses. For all three stimulus types, one direction from a set of 8 was selected in pseudo-random order, and the presentation of NCS, DBC, and LDG stimuli was also in a pseudo-randomized order.

### Mouse V1 single units respond to NCS stimuli

We analyzed the stimulus-evoked spiking of 520 single units. Example responses evoked by the NCS, LDG, and DBC stimuli are shown in Fig. 2a, b. We found that 57.2% of the units responded to both NCS and LDG stimuli, 39.5% responded to only LDG stimuli, and 3.3% responded to only NCS stimuli. In contrast, the DBC stimuli did not evoke any response (Fig. 2c). The magnitude of the response to NCS stimuli was significantly smaller than the response to LDG stimuli (Wilcoxon rank-sum test: Z=12.2, p=2.9e-34) and there was no evoked response to the DBC stimulus despite its exact pixel-wise changes compared to the NCS stimulus (Fig. 2d, e, f). Importantly, the lack of response to the DBC stimuli demonstrates that the NCS response is not due to the local physical changes of stimulus.

**Fig. 2:**
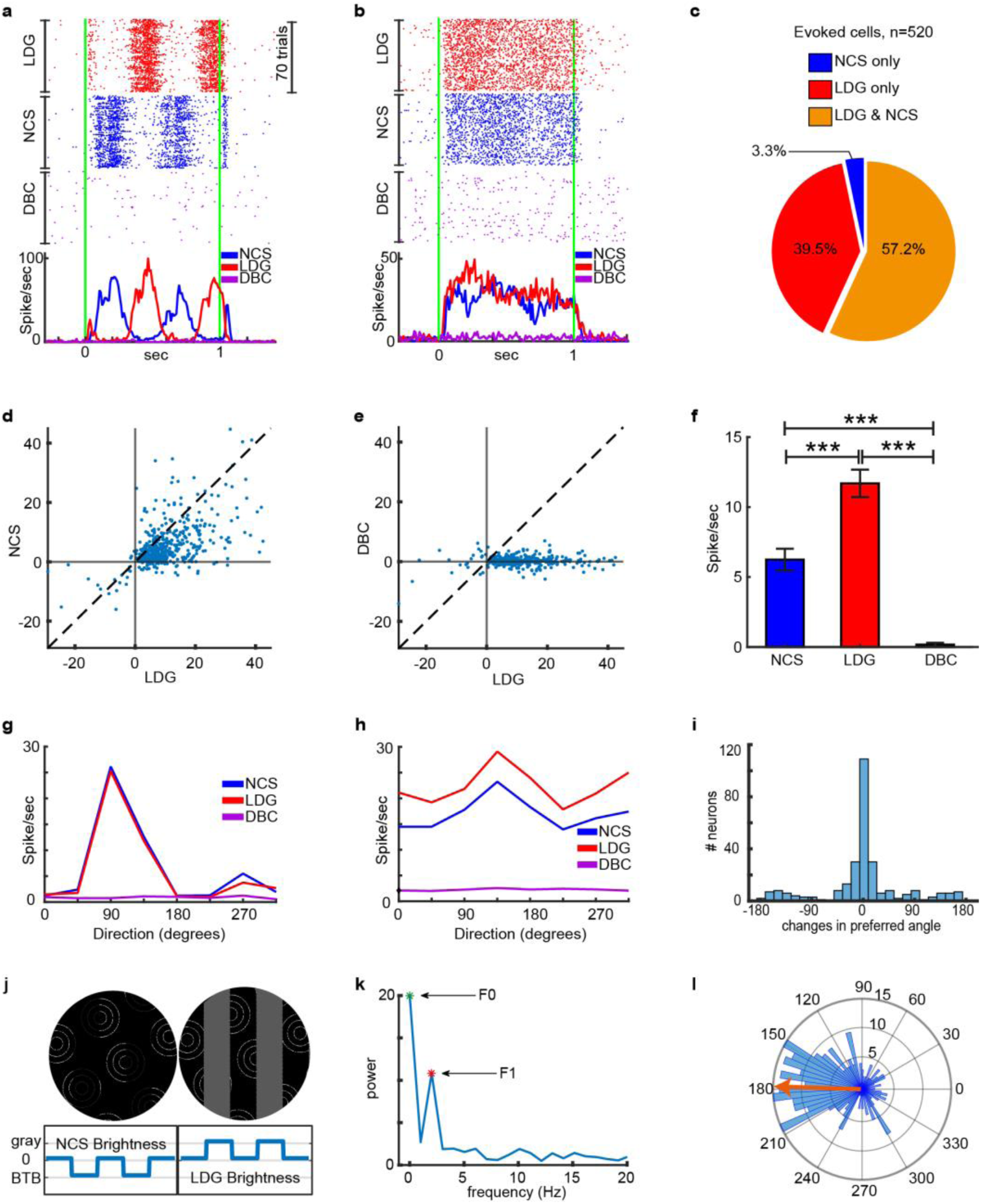
Single units respond to NCS stimuli with tuning properties characteristic of a response to a physical grating. **a and b**, Rasters and peri-stimulus time histograms (baseline-subtracted) of two V1 units in response to different stimulus types. The plots show the response to physical gratings (LDG stimuli), illusory gratings (NCS stimuli), and diffusion blocked illusory gratings (DBC stimuli) presented at the preferred direction of each unit (90° and 135°, respectively). The green lines indicate the times of stimulus onset and offset. Unit a responses are in accord with simple cell properties that phase-lock to grating stimuli, whereas responses of unit b are phase-independent and correspond to complex cell behavior. **c**, The pie chart shows the percent of single units that responded to the various stimuli. **d**, The scatter plot illustrates the maximal (trial-averaged) firing rate (spike/sec) of all 520 visually responsive units for NCS stimuli against LDG stimuli at the preferred orientation angle of each unit. **e**, The scatter plot shows the maximal response of all 520 units for LDG and DBC stimuli. Plotting conventions are identical to panel d. **f**, The bar plot shows the average response magnitude across all 3 stimulus types (N=520 units). Error bars indicate 95% confidence intervals. *** indicates p<0.0001. **g and h**, The direction tuning curves of the two example single units shown in panels a and b. **i**, The histogram shows the differences in the preferred angle between responses to NCS and LDG stimuli for the 297 units (57.2% of 520 units) that responded to both stimulus types. **j**, Due to the location of the grey segments in the NCS stimuli, the illusory grating is 180° out of phase with the physical grating (LDG stimuli). Example stimuli and a schematic of grating bar brightness are shown to illustrate that the dark bars of the illusory grating are aligned with the light bars of the physical grating. **k**, Frequency profile of single unit with an F1 dominant component of the power spectrum. **l**, The polar plot shows the phase shift between the response evoked by NCS and LDG stimuli (N = 209 units with an F1 dominant component in the evoked response). The red arrow shows the angular mean of this circular distribution (178.2°). The radial numbers indicate the number of neurons in the histogram.

As primary visual cortex neurons exhibit preferred angles for drifting gratings across species including mice^35-37^, we reasoned that if mouse V1 single units respond to illusory gratings as though they are perceived like gratings that are physically present, then the preferred angle would be similar for physical and illusory gratings. We obtained the preferred angle of each single unit using the eight drift directions of the LDG and NCS stimuli. The preferred angle was defined as the drift direction that evoked the maximal response for each unit. We found that the preferred angle was invariant for most single units when comparing the LDG and NCS stimuli (Fig. 2g, h, i). Therefore, for any given preferred angle determined by physical gratings, the single unit tended to prefer the same angle when illusory gratings were evoked by NCS stimuli.

Given that humans perceive the illusory grating as darker than the surrounding black background (Fig. la, Supplementary Movie 1), we presented NCS and LDG stimuli with a 180° relative luminance phase shift (Fig. 2j) in order to demonstrate that mouse V1 units respond to the illusory grating as if the bars are darker than the surrounding black background. Such a result would strongly support that mouse V1 units respond to illusory brightness. In order to demonstrate this property in V1 units, we tested the hypothesis that unit responses preserved the spatial phase properties of the stimuli. As is shown in raster and peri-stimulus time histograms (PSTHs) of example units, the response to NCS stimuli is shifted compared to the response to LDG stimuli (Fig. 2a, b). We quantified this effect by calculating the phase shift between the first harmonic (Fl component) of the neuronal responses to NCS and LDG stimuli. In order to ensure the reliability of the calculated phase, this analysis was only applied to a subset of units (N = 209) in which their Fl component was the dominant frequency component (i.e., the power of Fl component corresponding to the 2 Hz temporal frequency of the grating is larger than all other non-zero components). An example of an Fl dominant unit is shown in Fig. 2k. We found that the phase shift between the NCS response and LDG response was significantly non-uniform (Rayleigh’s test, Z=50, p=7.3c-24) and tightly distributed around a circular mean of 178.23° (95% confidence interval = [167.66°, 188.90°]). This 180° phase shift in unit responses between NCS and LDG stimuli indicates that mouse V1 units respond to illusory brightness in the form of grating bars that are darker than the surrounding black background. Overall, the results presented in Fig. 2 indicate that mouse V1 units respond to the illusory gratings evoked by NCS stimuli in a fundamentally similar manner to how they respond to physically present gratings.

### NCS responsive units are at a higher level in the V1 functional hierarchy

We next aimed to determine at what level the NCS stimulus-evoked response, and thus neural processing of the brightness illusion, fell in the local cellular hierarchy within VI. Response latency has been considered as an independent measure of the hierarchical level of V1 neurons^38^. Moreover, studies on humans^39^, macaque^40^, and mice^41^ have found that the neuronal response to visual illusions is delayed compared to physical stimuli, which may suggest that the neural correlates of visual illusions may be at a different level within the V1 cellular hierarchy. For instance, a later response to a stimulus might be due to additional serial synaptic interactions both within V1 and from top-down inter-cortical poly-synaptic inputs from higher levels of the cortical hierarchy that may contribute to illusory perception^38-41^ We calculated the latency of the response evoked by the different stimuli using only stimuli at the preferred direction of each unit^42^ (see Methods section for a description of the method of calculating latency). We excluded simple cells, which have a phase-locked response to the drifting grating, and for which it is not possible to estimate response latency. We found that NSC stimulus-evoked responses are later than responses to LDG stimuli (Fig. 3a). The mean ± SEM of latency was 65.74±0.17 ms for LDG but increased to 99.18±0.33 ms for NCS stimuli (Wilcoxon rank-sum test: Z=7.43, p=l .06e-l3). This delay in response time for the illusory stimulus may indicate that additional serial synaptic activations are required prior to the activation of V1 neurons. These results demonstrate that the V1 neuronal response to illusory gratings is delayed relative to physical gratings and may indicate that its neuronal correlate is at a different level of the visual hierarchy.

**Fig. 3:**
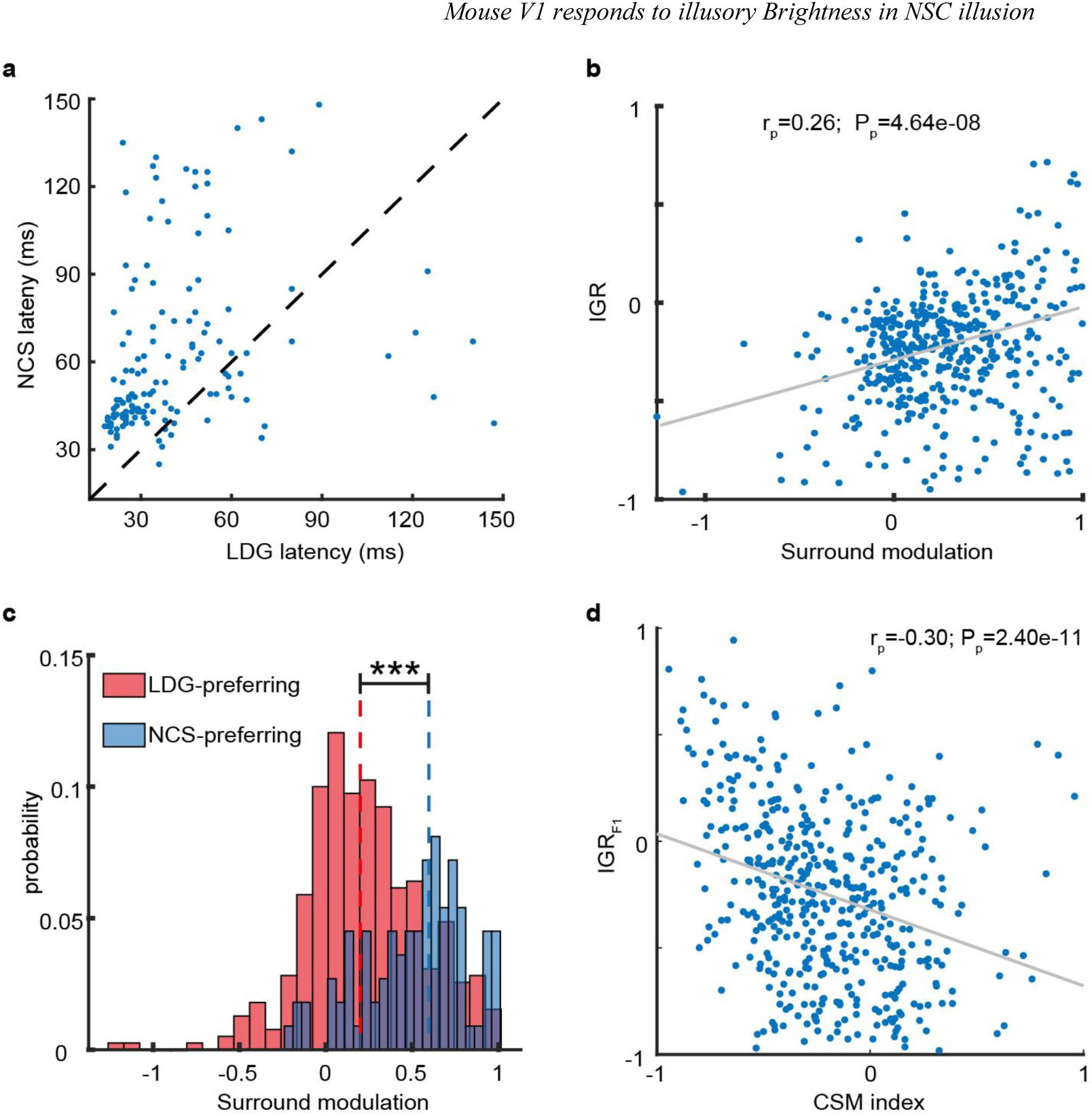
NCS preferring units respond earlier, receive more surround modulation, and may correspond to complex cells. **a**, The response latencies to NCS and LDG stimuli. Points above the dotted line indicate a later response to NCS stimuli relative to LDG stimuli. Each point represents a single unit. **b**, IGR plotted against surround modulation index. The fit line and rho value and p-value for Pearson’s correlation coefficient are shown. **c**, Distributions of surround modulation index plotted separately for NCS-preferring units (with positive IGR) and LDG-preferring units (with negative IGR). ^***^ indicates p<0.0001, Wilcoxon rank-sum test. **d**, The IGR_F1_ is plotted against complex-simple modulation index. Plotting conventions are as in panel b.

We further probed where V1 neuronal responses to illusions occur in the V1 cellular hierarchy by assessing the relationship between surround modulation and the NCS response magnitude for each unit. It has been suggested that surround modulation in V1 cells is a result of intra-V1 horizontal connections^43-46^ as well as inter-cortical feedback/feedforward connections^24,26^. Neurons with greater local horizontal inputs and/or top-down inter-cortical inputs can be considered as ‘higher level’ V1 neurons. Therefore, we hypothesized that V1 units with more robust surround modulation are preferentially responsive to NCS stimuli. We tested this hypothesis by calculating the correlation between an illusory grating response (IGR) index (see Methods section, Eq. 1) and a surround modulation index (see Methods section, Eq. 2). The IGR index quantifies the preference of units for the NCS stimuli relative to the LDG stimuli. A positive IGR indicates a greater preference for the NCS stimulus, whereas a negative value indicates a greater preference for the LDG stimulus. The surround modulation index was calculated using the size tuning curves of each unit. This index is negative when stimulus presentation outside the classical receptive field facilitates the firing rate (i.e., a so-called ‘facilitative cell’) and is positive when extra- classical receptive field stimulation suppresses the firing rate (i.e., ‘suppressive cell’). Supplementary Fig. 1 shows the size tuning of example facilitative and suppressive cells. We found that IGR and the surround modulation index were positively correlated (Pearson coefficient: r=0.26, p=4.6c-8). The relationship between these variables is shown in Fig. 3b. This result indicates that a greater response to NCS stimuli (relative to LDG stimuli) is associated with positive surround modulation indicative of suppressive cell activation. When we separated NCS preferring units and LDG preferring units, we found that LDG preferring units had a mean±SEM surround modulation index of 0.22±0.01, indicating weak surround modulation (Fig. 3c). On the other hand, for NCS preferring units, the mean±SEM was 0.53±0.03. The difference in surround modulation index between these unit sub-populations was significant (Wilcoxon rank-sum test: Z=7.15, p=8.4c-13). These results indicate that NCS preferring units have more robust surround modulation in comparison to LDG preferring units and that illusory brightness processing may involve VI suppressive cells. These findings provide evidence supporting the notion that VI units which respond to illusory brightness receive greater intra-V1 horizontal connections and/or inter-cortical connections and are therefore positioned at a higher level in the VI cellular hierarchy.

Another property of V1 cells related to functional hierarchy is their complex versus simple cell designation^29-32^. Unlike simple cells, there are no segregated excitatory/inhibitory areas in the RF of complex cells, and therefore their responses are not locked to the phase of the drifting grating. The response of two complex cells is shown in Supplementary Fig. 2. Many studies have suggested that complex cells receive convergent inputs from simple cells^22,23,31,47,48^. Thus, complex cells might be at a higher hierarchical level in VI. We, therefore, tested the hypothesis that NCS preferring units would be more likely characterized as complex cells. We separated putative simple and complex cells using a complex-simple modulation (CSM) index, which calculates the power of the Fl component of the grating-entrained response relative to the FO component (see Methods section, Eq. 3). The CSM index was calculated using responses to LDG stimuli. A CSM index of -1 is indicative of a complex cell, whereas an index of +1 indicates a simple cell. We observed no correlation (Pearson’s correlation, r=0.01, p=0.8) between CSM index and IGR (Supplementary Fig. 3). We next calculated the IGR index for NCS stimuli relative to LDG stimuli but used the amplitude of the Fl component rather than the average stimulus-evoked firing rate. The amplitude of the Fl component (calculated from the power spectrum of the evoked response) quantifies the degree of firing entrained to the 2 Hz temporal frequency of the grating. We refer to this version of the IGR index that compares the Fl amplitude for NCS versus LDG stimuli as the IGR_F1_ index (see Methods, Eq. 4). Fig. 3e shows that the IGR_F1_ index was negatively correlated with the CSM index (Pearson’s correlation, r=-0.30, p=2.4c-ll). This result suggests that units, which are likely to be complex cells, have a larger response entrained to the temporal frequency of the grating for NCS stimuli (an illusory grating) relative to the LDG stimuli (a physically present grating). These data provide additional support for V1 units that respond to NCS stimuli being in a higher position in the V1 cellular hierarchy.

### Putative inhibitory interneurons have a larger preferred response to the NCS stimulus than putative excitatory neurons

As interneurons in V1 receive more top-down inter-cortical input compared to principal neurons^25,33,34^, and given that top-down modulation of V1 is thought to play a role in the perception of illusions^41,49^, we hypothesized that V1 interneurons have differential responses to NCS stimuli. We used extracellular waveform characteristics to identify putative V1 interneurons and pyramidal neurons and compare their IGR and IGR_F1_ indices. We determined a putative neuron type using the trough-to-peak latency (TPL) for the average spike waveform of each unit^36,50,51^. The distribution of TPL values was bimodal, thus suggesting two classes of neurons: those with a narrow waveform were considered to be putative inhibitory interneurons (I units, N=146), and those with a wide waveform were putatively pyramidal neurons (E units, N=374). Fig. 4a shows the TPL distribution and separation of putative classes of neurons. LDG stimuli evoked responses in a similar proportion of I units and E units (95% and 92%, respectively; Fig. 4b). Importantly, however, NCS stimuli evoked responses in a larger proportion of I units compared to E units (69% and 57%, respectively; chi-squared test: χ^2^ =6.28, p=0.012). Moreover, the IGR was larger for I units relative to E units (Fig. 4c; Wilcoxon rank-sum test: Z=-3.5, p=4.3e-4). We also found that the IGR_F1_ index of I units was higher than that of the E units (Fig. 4d; Wilcoxon rank-sum test: Z=-3.49, p=4.7e-4). These results indicate a potential role of putative inhibitory interneurons in the neural response to illusory gratings evoked by the NCS stimuli.

**Fig. 4:**
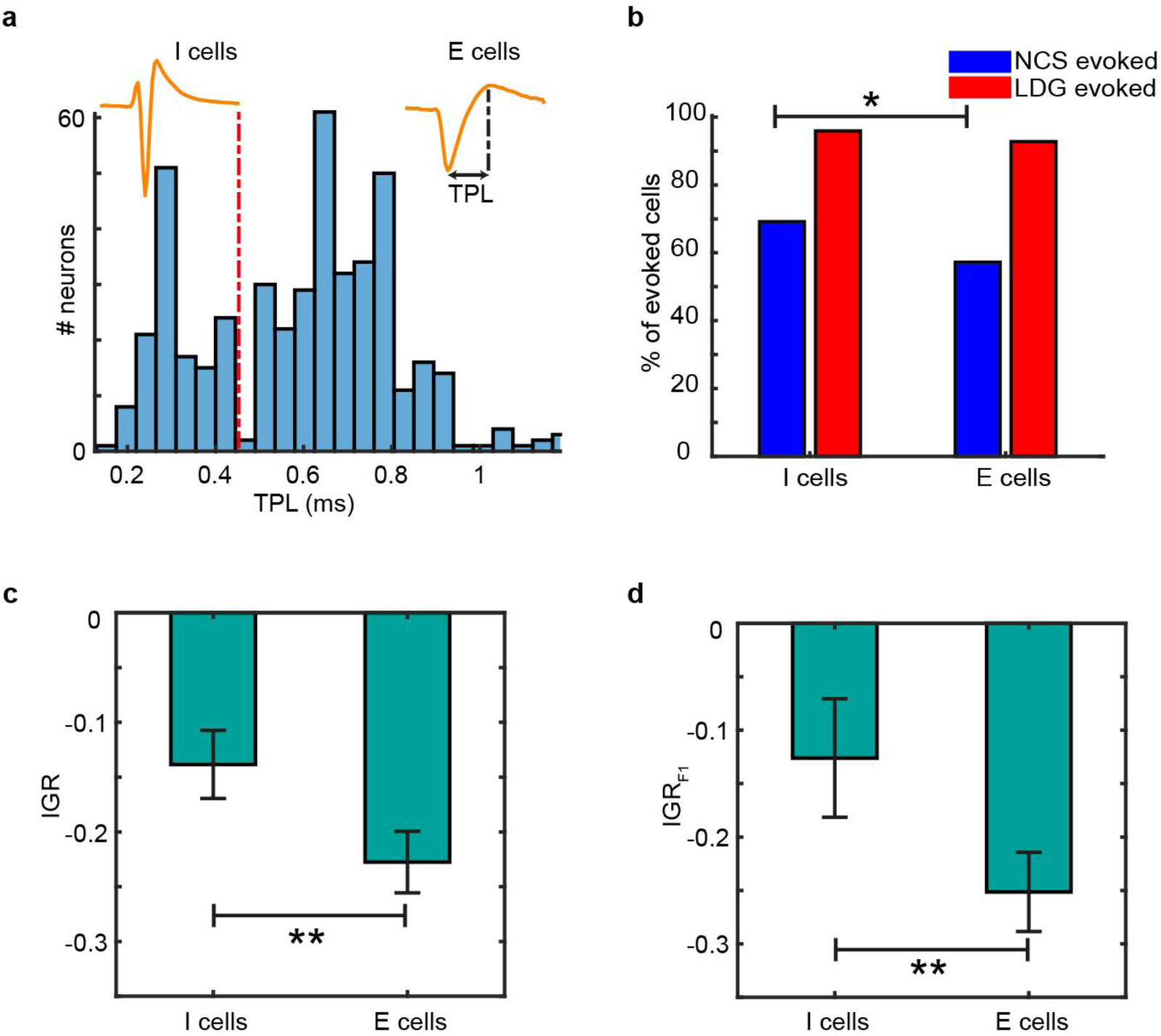
Putative inhibitory interneurons were preferentially responsive to NCS stimuli. **a**, The distribution of extracellular waveform TPL values for all recorded units. The dashed red line on the bimodal distribution shows the intersection of two distinct Gaussian distributions. Example spike waveforms are shown for one example unit from each of these distributions. Those with short TPL are putative interneurons and those with long TPL are putative pyramidal neurons. **b**, The percent of E units and I units with visually evoked responses to NCS and LDG stimuli. ^*^ indicates p<0.05, chi-squared test. **c and d**, The mean IGR and IGR_F1_magnitude of I units and E units (error bar = 95% Confidence Interval). ^**^ indicates p< 0.0005 Wilcoxon rank-sum test.

## Discussion

While the neural correlates of illusory contours has been extensively studied^40,41 52-55^, the neural correlates of illusory brightness have not been studied at the level of single neurons in any species. fMRI studies of human V1 BOLD signal responses to different features of illusory surfaces, specifically illusory fill-in in NCS9 and the Cornsweet illusion^10^, have been unable to discriminate between the perceptual experience of brightness, color, or other aspects of stimulus^12,14,49^. Moreover, only limited extrapolation can be made about neuronal activity from the fMRI BOLD signal. Although some single neuron studies have reported V1 cells that respond to surface luminance in Monkey^5^ and cat V1^56,57^, it is controversial whether V1 contributes to the perception of illusory brightness^8,16-18^, and thus, the neural correlates of illusory brightness remain unknown.

Here, we took advantage of NCS - previously only demonstrated in humans^9,21^ − to produce a brightness illusion that forms an oriented grating. We used NCS stimuli to probe the neural correlates of illusory brightness in mice. The specific use of gratings allowed us to compare multiple properties of the neuronal response between illusory and physical gratings, including drift direction (angle) tuning properties, entrainment to the temporal properties of the grating (i.e., the Fl component of the neuronal response), and phase shifting to illusory and physical gratings with opposing phase. Using this design, we show that mouse V1 single units respond to the illusory drifting grating evoked by NCS stimuli yet, critically, did not respond to control stimuli in which pixel-wise changes in physical luminance are matched to the NCS condition but illusory brightness is blocked. These control stimuli do not evoke the perception of illusory brightness or illusory gratings by human observers (see Supplementary Movie 2). We found that the neuronal tuning properties are similar for physically present gratings (LDG stimuli) and illusory gratings, which suggests that NCS stimuli evoke neuronal responses characteristic of those to actual gratings. Importantly, by presenting illusory gratings and physical gratings with a 180° spatial phase shift, we show that V1 neurons respond to the spatial phase properties of the illusory gratings and, therefore, track the illusory brightness perceived by human subjects. Collectively, these results are strong evidence for the response of V1 single units to illusory brightness.

The neuronal correlates of illusory perception are thought to depend on late-stage synaptic feedback to V1 neurons that occurs after an initial feed-forward pass through V1^41,49^. One prediction of this model is that the V1 neuronal response latency should be delayed for illusory stimuli relative to physical stimuli because of the time required for activation of additional synapses. Indeed, in line with the prior work in humans^39^, macaques^40^, and mice^41^, we found that the neuronal response to the illusory grating was delayed relative to a physical grating. Our findings support the notion that the V1 neuronal response to illusory gratings cannot be only feed-forward and driven by physical changes in the receptive field but also involve interactions among neurons within V1 and outside VI.

Surround modulation may be used as an indicator for activation of specific intra-V1 or extra-V1 neuronal connections. We found that V1 neurons with greater surround suppression effects have a greater response to NCS stimuli. This finding may support the notion that top-down feedback is involved in the V1 responses to NCS stimuli and illusory brightness perception because optogenetic studies have shown that surround suppression in V1 relies on intra-V1 horizontal connections^45^ and on feedback connections^25,58^. On the other hand, several studies have demonstrated surround suppression in the lateral geniculate nucleus (LGN), suggesting surround suppression could be partially inherited from LGN through feed-forward connections^59,62^. Although the intra-Vl and extra-Vl neuronal connections involved in illusory brightness processing remain unclear, our work shows a correlation with surround modulation that can be used to infer which neuronal connections are involved as the neuronal basis of surround modulation becomes better characterized.

We also evaluated the propensity for complex cells to respond to illusory gratings, given that their preferential activation by illusions may indicate the engagement of top-down feedback from outside VI. Complex cells receive input from simple cells within V1 and other visual areas^22,23,29-32,47,48^, and their activation is therefore dependent upon V1 horizontal connections and feedback connections. We found that complex cells had a higher Fl response to the illusory grating, which can be considered as evidence for their response being driven by feedback from higher visual areas. Note here that, while these results appear to be self-contradictory to the very definition of complex cells, where they have smaller Fl components than simple cells, we have calculated the simple-complex index using the evoked response to luminance-defined gratings. This finding is in line with a prior neuroimaging study in humans, which suggested that extrinsic inputs to V1 contribute to the neural correlates of filling-in in NCS^9^.

Finally, top-down cortical modulation also preferentially targets GABAergic interneurons in V1^25,33,34^ and thus, interneurons may play a crucial role in feature selectivity and visual perception^63-69^. We assessed cell type-specific responses to illusory gratings and found that putative interneurons were more responsive to NCS stimuli than LDG stimuli. This finding suggests that mouse V1 interneurons may contribute to the processing of illusory brightness, potentially because they are targeted by top-down modulation.

A final consideration concerns the relation between neural responses observed here and coding of the high-level percept of fore- and background. In contrast to luminance, which was phase-shifted by 180 degrees between NCS and LDG, the perceived fore-grounds had zero phase-shift. For responses reflecting the high-level percept, one could hence expect zero phaseshift. This would be mediated by so-called border-ownership selectivity (BOS), where the neural response is modulated according to side of the edge that constitutes the fore-ground^70^. However, this would be expected only for a small fraction of neurons, for two reasons. First, in primate VI, only about 20% of neurons show BOS. Second, nearly all of these BOS neurons are also luminance-polarity selective, and BOS does not necessarily override polarity response. Only about about 3% of neurons show only BOS^70^, which may correspond to the small fraction observed here with zero phase-shift. Similar to the characteristics observed in NCS responses here, also BOS occurs in well-connected neurons and within a short temporal delay compatible with a feedback modulation^71^.

In summary, we show that mouse V1 single units respond to illusory brightness evoked by NCS stimuli, which demonstrates that individual V1 neurons contribute to processing of illusory brightness. The illusory response in V1 is more robust in putative complex cells, putative inhibitory interneurons, and units with greater surround suppression, which supports the potential contribution of higher visual hierarchies in the neural correlates of illusory processing in neon color spreading.

## Acknowledgements

This work was funded by the Max Planck Society. This paper was typeset with the bioRxiv word template by @Chrelli: www.github.com/chrelli/bio-Rxiv-word-template.

## Author contributions

Conceptualization - MW; Data acquisition and curation - AS, KW, GN; Formal analysis - AS, MW, NT, AB; Methodology - AS, MW, NT; Project administration - MW; Supervision - MW, NT, AB; Visualization - AS, MW, NT; Writing - AS, MW, NT, AB; Resources - NKL.

## Data availability

Raw data is available upon reasonable request.

## Materials and Methods

### Subjects

Experiments were performed on head-fixed adult mice on a disc. The local authorities approved all animal procedures and in compliance with EU Directive 2010/63/EU (European Community Guidelines for the Care and Use of Laboratory Animals). Data acquisition was done through 32 electrode penetrations in both hemispheres of two C57BL/6 mice (male) and four PV-Cre mice (three males, one female; homozygous for the PV-Cre genes, B6;129P2-Pvalbtml(cre)Arbr/J).

### Surgical Preparation

Mice were induced by 2.5 % of isoflurane during surgery and maintained at 1-2 %. Also, Atropine (Atropinsulfat B. Braun, 0.3mg/kg) and Buprenorphine (0.1 mg/kg) were administered via subcutaneous injections to reduce bronchial secretions and as analgesics, respectively. The scalp was sterilized and opened to expose the lambda and bregma sutures. A lightweight head-post was installed onto the skull using an adhesive primer and dental cement (OptiBond FL primer and adhesive, Kerr dental; Tetric EvoFlow dental cement, Ivoclar Vivadent). A small well was built around the exposed area using dental cement. Two silver wires were implanted between the dura and skull over the frontal lobe as ground references for extracellular recordings. Then, the skull was covered with Kwik-Cast (WPI). The post-surgery analgesic (Flunixin, 4mg/kg) continued to be administered every 12 hours for three days, and antibiotics (Baytril, 5mg/kg) were administered for five days. After recovery, animals were habituated to head-fixation and placed on a disc for three days (0.5 hours/day). On the fourth day, a small craniotomy (1 mm2) was drilled above the V1 at 2.5 mm laterally and 1.1 mm anterior of the transverse sinus^72^ under general anesthesia. Electrophysiological recordings were started one day after craniotomy surgery and continued on consecutive days for as long as the neuron isolation remained of high quality. The craniotomy was covered with Kwik-Cast after each recording.

### Electrophysiological Recordings

Mice were head-fixed on a disc and allowed to sit or run on it in a dark and electromagnetic isolated room. A 32-channel linear silicon probe (Neuronexus, Alx32-5mm-25-177-A32) was penetrated perpendicularly to -900 pm below the brain surface. Electrical signals were amplified and digitized at 30 kHz by the Cerebus data acquisition system (Blackrock Microsystems LLC) or RHD recording system (Intan Technologies). A photodiode was attached to the lower right corner of the screen to capture the exact stimulus onset from a white square synchronized to the stimulus presented.

### Visual Stimulation and Experiment Design

Stimuli were projected onto a gamma-corrected LED monitor (Dell U2412M, 24 inches, 60 Hz) placed 15 cm in front of the animal’s eye. Visual stimuli were programmed and generated in MATLAB (MathWorks, Inc.) and Psychophysics Toolbox Version 3 (PTB-3). To obtain the receptive field map of recorded neurons, black squares (15° widths) were presented on a gray background with a duration of 100 ms and an interval of 100 ms in different locations of 8 by 13 grids. The duration of the receptive field mapping session was 20 minutes. After this section, the response to each square was extracted by an analysis of multi-unit activity (MUA). Then the center of the MUA receptive field was estimated by the best fit of a two-dimensional Gaussian to the MUA activities. Subsequent target stimuli were presented on a gray background at the estimated receptive field center. To obtain the size-tuning curve, circular patches of drifting grating (spatial frequency 0.05 cycles/degree, temporal frequency 3 Hz) with different sizes (2.5, 5, 10, 15, …, 45 degrees) and two drifting directions (rightward and upward) were presented. The duration of the stimulus presentation was 666.7 ms with a 500 ms interval, and the duration of the whole session was 25 minutes.

We presented three types of drifting grating stimuli for the neon color session. (1) Neon color spreading (NCS) consists of nine patches of white concentric circles (0.1° thickness) as inducers (each patch had three circles), arranged on a three-by-three virtual grid on a black square (35° widths). The diameter of the inducers was 3,6, and 9 degrees, respectively. At each frame, the intersection of concentric circles and a drifting grating (spatial frequency =0.05 cycle/degree and temporal frequency= 2 Hz) was replaced with gray segments, resulting in the “darker than black” illusory grating (Fig. la, Supplementary Movie 1). (2) Diffusion-blocked stimulus is a control condition with exactly the same pixel-wise changes as NCS, while each inducer circle is sandwiched by two white circles (0.4 thickness). The added circles constrain the gray filling-in and reduce the illusory effect (Fig. 1c, Supplementary Movie 2). (3) Luminance-defined grating (LDG) is defined as gray grating moving on top of inducers (Fig. Id, Supplementary Movie 3). These three types of drifting grating were generated in eight directions, making 24 conditions. These stimuli were presented with a duration of 1 second for 70 trials in a pseudo-randomized order.

### Data analysis

For spike detection and clustering, we first concatenated the recorded data in all three experiment sessions (i.e., receptive field mapping, size tuning, and neon color). We then used the Kilosort algorithm, a template matching algorithm written in MATLAB for spike sorting, with the default parameters^73^. A manual clustering followed this for further merging, splitting, and choosing isolated clusters using template-gui74. All further analyses were done in MATLAB using built-in functions and CircStat Toolbox^75^. The peristimulus time histogram (PSTH) initially was calculated with a resolution of 1 ms and smoothed by a moving average window of 2 ms. To extract visually evoked neurons and estimate their onset latency, we assumed that the spontaneous spiking activity prior to stimulus presentation follows a Poisson distribution^42^. By fitting a Poisson distribution to 300 ms prior to stimulus onset, the spontaneous firing rate À was estimated. If the spiking activity after stimulus onset deviates from the background Poisson distribution to a particular level in three consecutive bins (a probability of p< 0.01 for the first two bins and p< 0.05 for the third bin), the neuron was considered as an evoked neuron, and the corresponding time for the first bin is considered as a response latency of the neuron42. The preferred angle of cells is defined as the stimulus direction with the maximum response. To calculate changes in the preferred angle of neurons in NCS and LDG, first, we captured the direction tuning curve of the neuron by taking the average response of cells during the stimulus presentation (one second). Then we interpolated the tuning curve with the spline method to get a more precise estimation of the preferred angle.

We have quantified the illusory grating response for each neuron as follows.

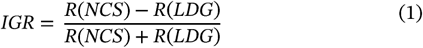

Where R is the average response of the neuron to the stimulus over the presentation period. A positive IGR represents a higher response to NCS and vice versa.

We defined the surround modulation index as

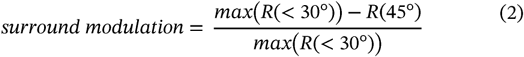

The surround modulation index is negative for facilitative cells and positive for suppressive cells.

We implemented Fast Fourier Transform (FFT) on the PSTH to extract a temporal component of neuronal responses. The Fl component of the response is the power for 2 Hz, which is the same as the temporal frequency of the stimulus, and the F0 component is the average response.

The complex-simple modulation index was calculated using the following equation for every single unit:

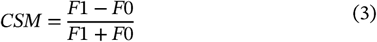

Where Fl is the power of 2 Hz frequency, and F0 is the average firing rate in response to LDG stimuli. Neurons with a positive CSM index have a phase-locked response to the temporal frequency of drifting grating and are classified as simple cells. The phase-locked response is due to the separated excitatory and inhibitory subregions in their receptive field. A negative CSM index indicates the degree of spatial invariance in the receptive field and a lack of phase-locked responses.

We calculated the relative amplitude of the Fl component (IGR_F1_) as in the following,

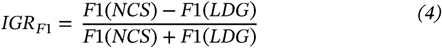

where Fl(NCS) and Fl(LDG) are the amplitude of the Fl component of the response to NCS and LDG respectively.

To classify cells into two groups of putative inhibitory and excitatory cells, we fitted two Gaussian functions to the histogram of TPL, which shows a bimodal distribution. The intersection point of two Gaussian curves was selected as a threshold to classify putative I/E cells (Supplementary Fig. 4). We have used the two-sided Wilcoxon rank-sum test and Chi-squared test for the statistical analysis, and for directional statistics, Rayleigh’s test has been implemented.

## Supplementary information

**Supplementary Fig. 1:**
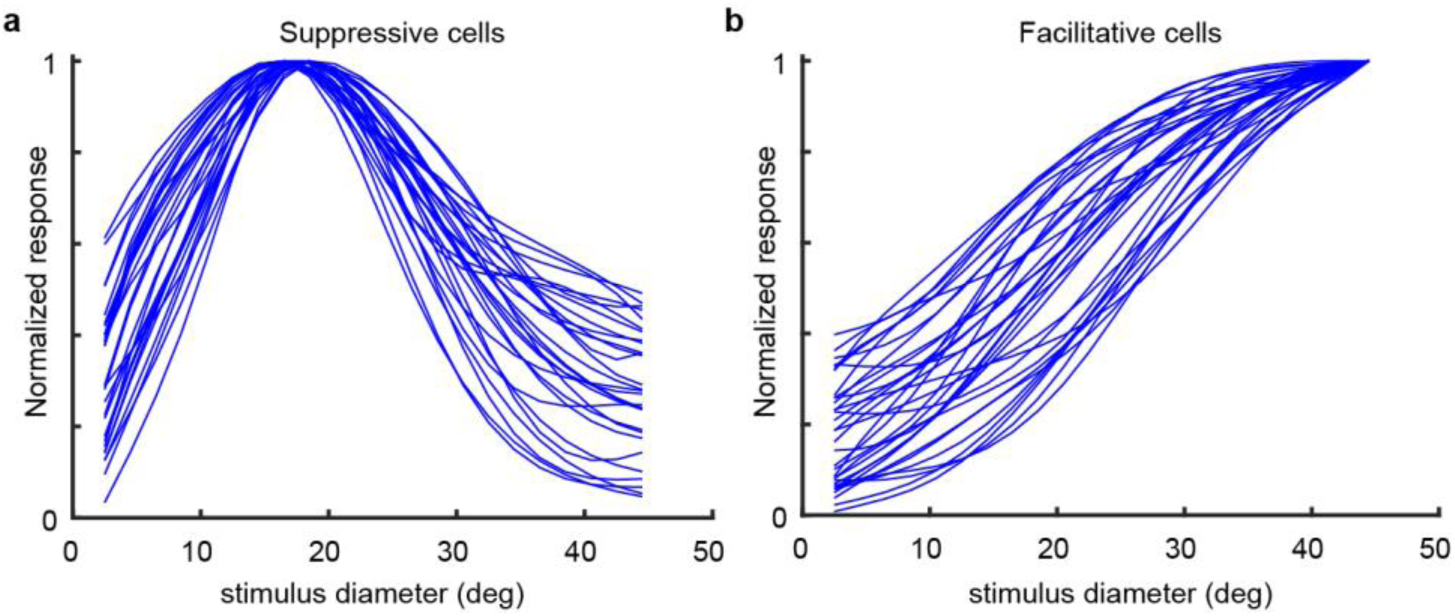
Suppressive and facilitative cells. **a**, size tuning curve of 30 example of suppressive cells. **b**, size tuning curve of 30 example facilitative cells.

**Supplementary Fig. 2.**
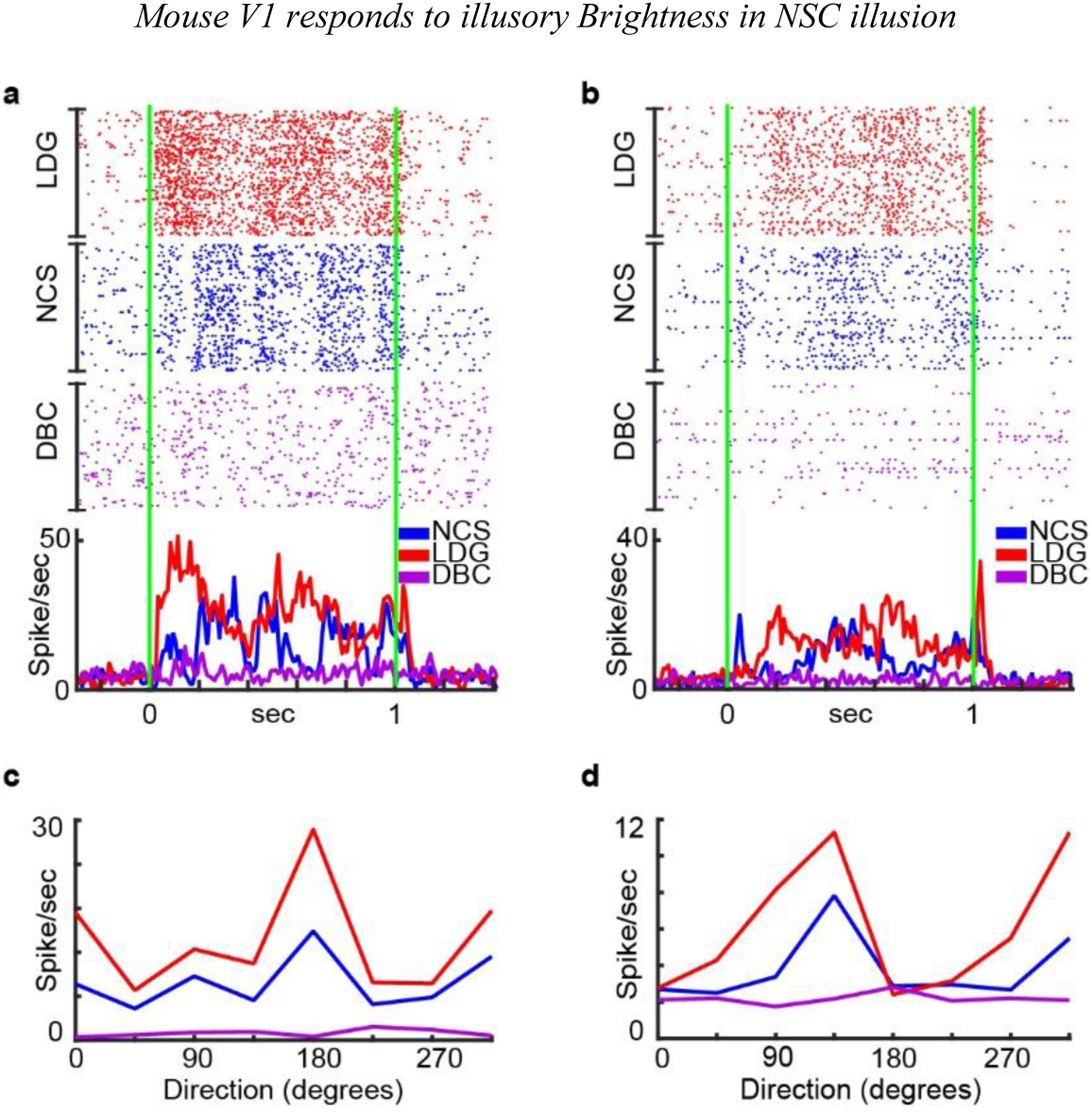
Example of complex cell responses. **a and b**, Raster and peri-stimulus time histogram (baseline-subtracted) of two V1 complex cells (CSM in a: -0.7 and in b:-0.6) in response to different conditions at the preferred direction of the neuron (180° and 135°). Green lines show the stimulus onset and offset. **c and d**, direction tuning curves of the two example neurons in **a** and **b**

**Supplementary Fig. 3.**
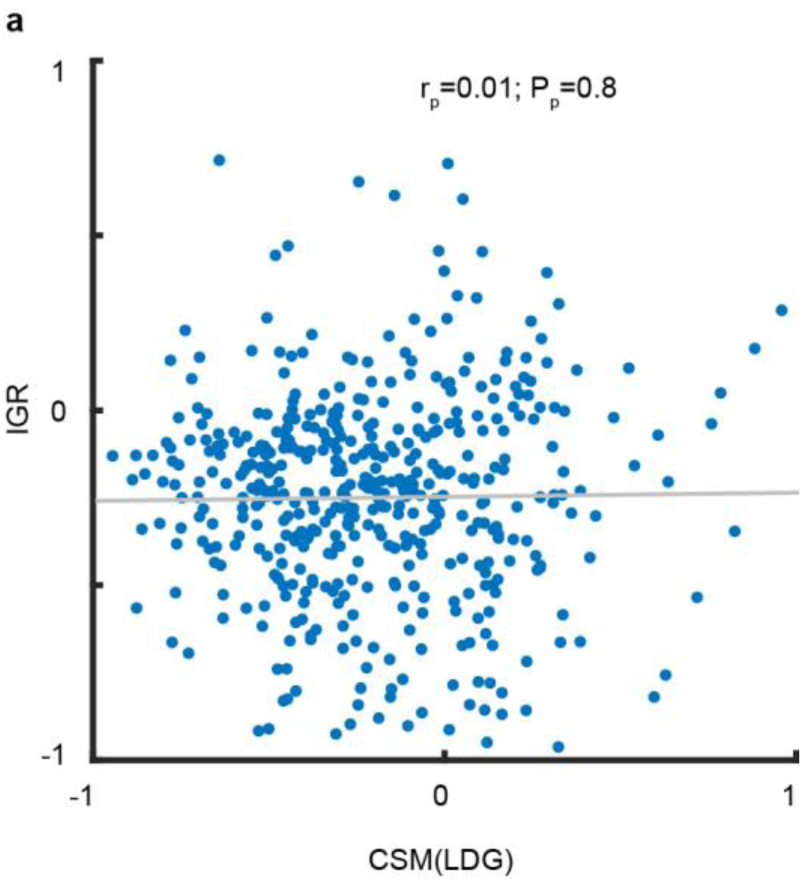
Illusory grating response plotted against complex simple modulation (CMS). There is no significant correlation (r=0.01, p=0.8) between CSM and IGR.

**Supplementary Fig. 4.**
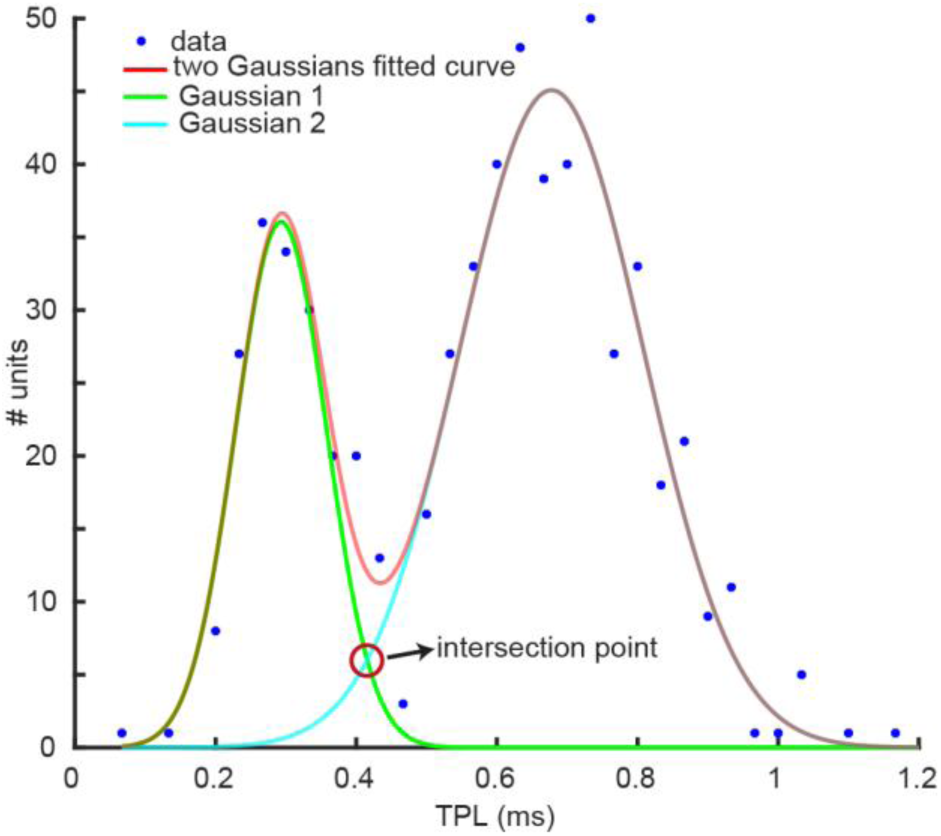
Classification of putative I/E cells. Two Gaussian functions are fitted to the distribution of trough-to-peak latency (TPL), and the intersection point is selected as a threshold. Cells with a TPL shorter than the threshold are considered as narrow-waveform inhibitory cells. Other cells are regarded as wide-waveform principal cells (i.e. putative excitatory cells).

**Supplementary Movie 1**.

Neon color spreading (NCS) stimuli with different directions. (ATTACHED)

**Supplementary Movie 2**.

Diffusion-blocked control (DBC) stimuli with different directions. (ATTACHED)

**Supplementary Movie 3**.

Luminance defined grating (LDG) stimuli with different directions. (ATTACHED)

